# Reconsidering the Use of Dimethyl Sulfoxide for Xenobiotic-Gut Microbiota Interaction Studies

**DOI:** 10.64898/2026.08.12.743806

**Authors:** Qiwen Cheng, Hannah Glesener, Aurely Sanchez Carreon, Lee E Voth-Gaeddert, Rosa Krajmalnik-Brown

## Abstract

**Introduction:** Gut microbiota are vulnerable to foreign chemicals (xenobiotics) including pharmaceuticals, environmental pollutants, and dietary contaminants such as aflatoxin B1 (AFB1) and fumonisin B1 (FB1). Assessing the effect of these xenobiotics in the laboratory requires their dissolution in a solvent vehicle, such as dimethyl sulfoxide (DMSO). While DMSO is typically used at low concentrations under the assumption of neutrality, its independent impact on microbial dynamics is a potential experimental confounder that has not been fully explored.

**Methods:** Human fecal microbiota were cultivated *in vitro* for 16 days 16 days, supplemented with 0, 10, 100, and 1000 ppb of the tested xenobiotics (AFB1 or FB1) in 0.05% DMSO (v/v), with a DMSO-free control included for comparison. Microbial community dynamics were characterized via full-length 16S rRNA gene sequencing, and metabolic activity was assessed by measuring production of short-chain fatty acids and gases.

**Results:** DMSO significantly altered microbial metabolism and drove the consistent enrichment of *Desulfovibrio desulfuricans*. This shift occurred across all AFB1 and FB1 treatment groups regardless of their concentrations, indicating that the biological impact of the DMSO vehicle overshadowed the specific effects of the xenobiotics.

**Discussion:** These findings demonstrate that DMSO can induce significant microbial shifts independent of the xenobiotics under study, potentially confounding biological interpretations. This highlights a critical need for rigorous vehicle validation and the identification of safe thresholds for solvents used in microbiota research.

## 1. Introduction

The gastrointestinal (GI) tract serves as the primary interface between the host and the external environment, housing an intricate and densely populated ecosystem of microorganisms collectively termed the gut microbiota. This community, encompassing bacteria, archaea, viruses, and eukaryotes, is fundamental to the maintenance of host health. This complex ecosystem modulates various physiological processes, including the fermentation of non-digestible dietary fibers into short-chain fatty acids (SCFAs), which serve as primary energy sources for colonocytes and regulate systemic metabolism (Fan and Pedersen, 2021). Disruption in microbial fermentation can promote the translocation of pro-inflammatory microbial products, such as lipopolysaccharides, leading to metabolic endotoxemia and systemic inflammation. Furthermore, the microbiota play a critical role in the maturation and calibration of the host immune system, promoting the differentiation of regulatory T cells and maintaining the integrity of the intestinal barrier to prevent pathogen translocation (Round and Mazmanian, 2009; Belkaid and Hand, 2014). Beyond the gut, these microorganisms communicate with distant organs via the production of metabolites and neurotransmitters, influencing a broad spectrum of metabolic and neurological functions (Griffiths et al., 2026). For instance, the microbiota-gut-brain axis can function as a bidirectional communication network that links the GI tract and the central nervous system, producing neurotransmitters like gamma-aminobutyric acid and serotonin that modulate brain function and behavior (Mayer et al., 2022; Loh et al., 2024). Dysfunction in this axis serves as a hallmark of conditions such as Parkinson’s, autism spectrum disorder, and Alzheimer’s Disease (Sampson et al., 2016; Kang et al., 2017; Morton et al., 2023; Seo et al., 2023).

Given their residence at the primary site of nutrient assimilation and mucosal exposure, gut microbiota are inherently vulnerable to structural and functional shifts induced by foreign chemicals (i.e., xenobiotics). Xenobiotics encompass any synthetic or natural chemical compound, such as dietary components, environmental pollutants, industrial chemicals, and pharmaceuticals, that is foreign to a living organism’s biological system (Maurice et al., 2013; Koppel et al., 2017). While many are benign or therapeutic, exposure to toxic xenobiotics can trigger microbial dysbiosis and alter host metabolic networks. Furthermore, some compounds may exacerbate toxicity through downstream microbial transformations that convert otherwise benign precursors into highly toxic or carcinogenic secondary metabolites (Maurice et al., 2013; Koppel et al., 2017; Weersma et al., 2020). Among xenobiotics, foodborne mycotoxins—secondary metabolites produced by fungi—have emerged as a concern due to their ubiquitous presence in the global food supply and high toxicity (Liew and Mohd-Redzwan, 2018; Eskola et al., 2020). Specifically, aflatoxin B1 (AFB1) and fumonisin B1 (FB1) represent the most significant threats to public health due to their frequent occurrence in staple commodities like maize (Voth-Gaeddert et al., 2020). AFB1, a difuranocoumarin derivative produced primarily by *Aspergillus flavus* and *Aspergillus parasiticus*, is categorized by the International Agency for Research on Cancer as a Group 1 human carcinogen, primarily linked to the etiology of hepatocellular carcinoma (Chen et al., 2023; Choi et al., 2025). In parallel, FB1, an aminopolyol metabolite of *Fusarium verticillioides* and *Fusarium proliferatum*, is classified as a Group 2B carcinogen (Stockmann-Juvala and Savolainen, 2008; Chen et al., 2021; Anumudu et al., 2024). Evidence from animal studies suggests that the ingestion of AFB1 and FB1 induces gut microbiota dysbiosis, characterized by significant alterations in microbial composition, diversity, and metabolic profiles (Liew and Mohd-Redzwan, 2018; Guerre, 2020; Choi et al., 2025). Documented microbial shifts include changes in the abundance of beneficial taxa, such as *Lactobacillus* and *Bifidobacterium*, which play key roles in barrier integrity and immune modulation (Dang et al., 2017; Yang et al., 2017; Mateos et al., 2018; He et al., 2019; Zhang et al., 2021; Li et al., 2022; Liu et al., 2022; Guo et al., 2023; Ye et al., 2023, 2025; Chen et al., 2024). Meanwhile, exposure to these mycotoxins facilitates the expansion of opportunistic pathogens, such as Escherichia coli, which may exacerbate intestinal inflammation and promote metabolic dysfunction (Oswald et al., 2003; Voth-Gaeddert et al., 2019; Pu et al., 2021; Guo et al., 2023). These observations vary by host species, mycotoxin dosage, and exposure duration, yielding diverse microbial profiles across the literature. Conversely, microbial enzymes facilitate the sequestration or degradation of these mycotoxins, yielding metabolites characterized by reduced toxicity or diminished systemic bioavailability (Rushing and Selim, 2019; Guerre, 2020). For instance, certain lactic acid bacteria are known to bind mycotoxins to their cell walls, limiting their absorption across the intestinal epithelium (Choi et al., 2025).

Evaluating the interactions between xenobiotics and gut microbiota, *in vitro* or *in vivo*, often requires the use of solvent vehicles due to chemical hydrophobicity. While solvents such as methanol, ethanol, and acetonitrile are frequently employed, dimethyl sulfoxide (DMSO) remains the most common choice due to its superior solubilizing capacity. DMSO has been used to study xenobiotic-gut microbiota interactions at concentrations between 0.001% and 1% (v/v) (Chen et al., 2018; Lai et al., 2018; Tang et al., 2021; Li et al., 2024; Müller et al., 2024; Tian et al., 2024; Yang et al., 2024; Roux et al., 2025; Wang et al., 2025). However, the potential for DMSO to alter gut microbiota profiles remains poorly characterized, with many studies assuming non-inhibitory effects based on previous research. Our study reveals that DMSO could significantly shift human gut microbial metabolism and drive the enrichment of the sulfate-reducing bacterium *Desulfovibrio desulfuricans*. Notably, this microbial shift was consistent across all AFB1 and FB1 treatment groups regardless of their concentrations, indicating that the biological impact of the DMSO vehicle overshadowed any potential effects of the tested compounds themselves. Our findings suggest that DMSO may confound biological assessments by inducing microbial shifts independent of the xenobiotics being studied.

## 2. Materials and methods

### 2.1 Fecal sample collection

Fecal samples were obtained from 19 healthy volunteers in 2017. Participants were enrolled based on the following eligibility criteria: (1) Aged 18-65; (2) Affiliation with Arizona State University (students, faculty, or staff), or their friends and relatives; (3) No history of medical treatments within the previous two months; (4) No use of antibiotics, antifungals, or probiotic supplements (dietary yogurt consumption was permitted) within the previous three months; (5) No known systemic or chronic conditions (e.g. hepatitis or autoimmune diseases); (6) No current or past history of gastrointestinal diseases or persistent symptoms (e.g., Crohn’s disease, chronic diarrhea, or constipation); 7) Body Mass Index within the normal range. Exclusion criteria included individuals under 18 years of age, pregnant women, prisoners, individuals with physical or mental disorders, and those unable to provide informed consent. Fecal samples were self-collected by volunteers in sterile containers and immediately transferred to ice. Samples were then aliquoted in an anaerobic glovebox, preserved in a 50% (v/v) sterile glycerol solution and stored at -80 ℃ prior to use.

### 2.2 Stock solution preparation

Stock solutions of aflatoxin B1 (AFB1; Sigma-Aldrich) and fumonisin B1 (FB1; Cayman Chemical) were prepared using dimethyl sulfoxide (DMSO) as the solvent. A primary stock solution (2 µg/µL) was established by dissolving 5 mg of AFB1 or FB1 in 2.5 mL of DMSO. Subsequently, two additional stock solutions (0.2 and 0.02 µg/µL) were prepared via serial dilution of the primary stock with DMSO.

### 2.3 Bioreactor operation

The bioreactors were operated in two stages: an acclimation phase and an exposure phase (**Figure 1**). During the acclimation phase, fecal samples from the 19 volunteers were thawed at room temperature and homogenized in an anaerobic glovebox to create pooled feces. These feces were inoculated into quadruplicate 250 mL serum bottles containing 120 mL of growth medium. The bottles were sparged with nitrogen gas to remove oxygen, and incubated at 37 °C with shaking at 150 rpm for 48-hour cycles. At the end of each cycle, 80% of the medium was removed and centrifuged to harvest biomass, which was then transferred into fresh medium under anaerobic conditions to maintain growth. These transfers were repeated until the microbial community reached the steady state, defined as the point when 48-hour short-chain fatty acid (SCFA) concentrations and gas yields showed no significant variations between successive transfers (*p*> 0.05). Following acclimation, biomass from the quadruplicate bottles were pooled to generate a pooled culture. This was distributed into 28 mL anaerobic culture tubes containing 10 mL of medium supplemented with 10, 100, or 1000 ppb of AFB1 or FB1. A DMSO solvent control (i.e., 0 ppb mycotoxin) and a No-DMSO biotic control were included to account for solvent effects. An abiotic control with medium and DMSO was used to assess other possible chemical transformations or biological contamination. All treatments were conducted in triplicate. Consistent with the acclimation phase, 80% of the medium (8 mL) was removed from each tube every 48 hours and centrifuged to harvest the biomass. This collected biomass was then transferred into new tubes containing 10 mL of fresh medium with the corresponding mycotoxin concentration. The remaining 20% of the medium was reserved for SCFA analysis and DNA extraction. Comprehensive sampling was performed at the conclusion of 2-, 8-, and 16-day incubation periods, at which point headspace samples were collected for gas composition analysis, and liquid samples were harvested for SCFA characterization and DNA extraction.

**Figure 1.**
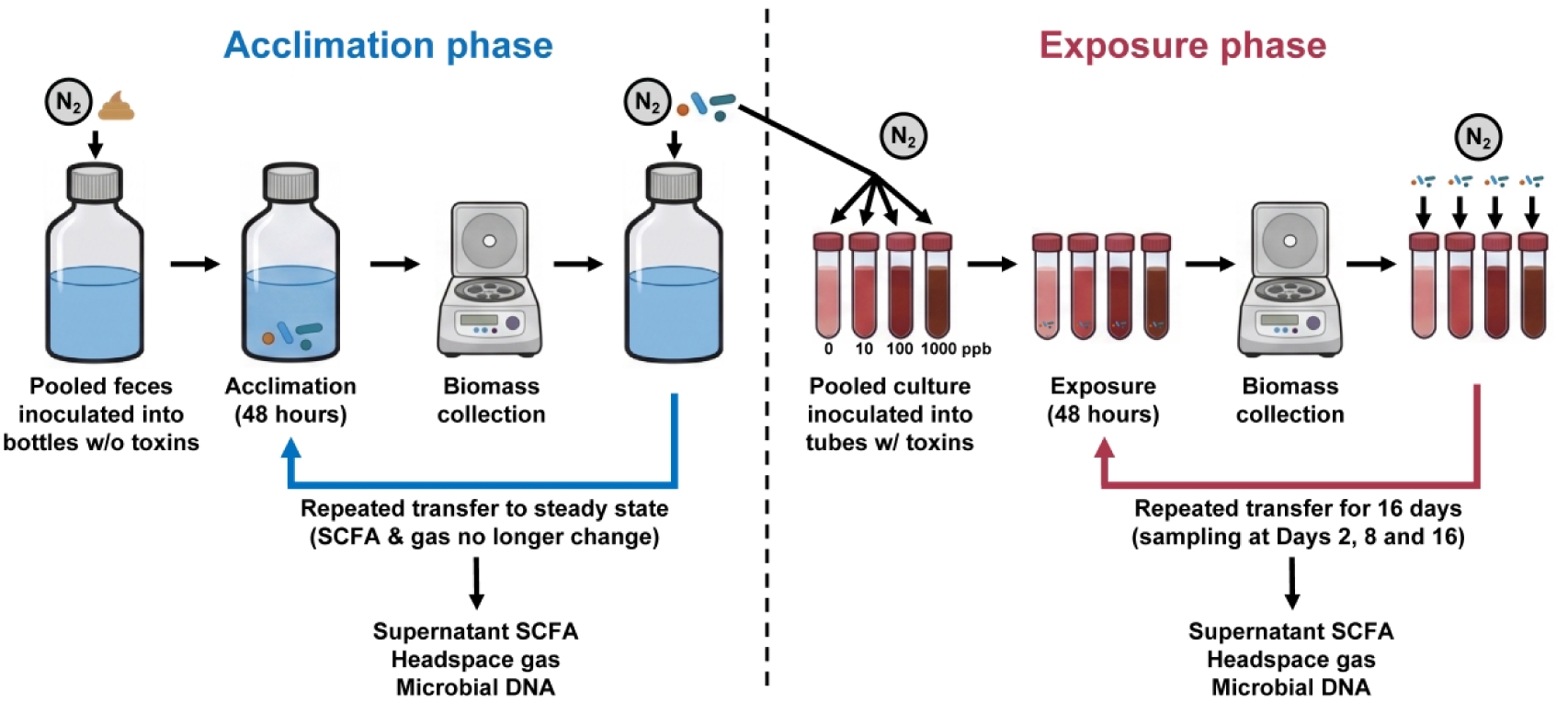
Schematic overview of the *in vitro* experimental design, including the acclimation and exposure phases. The workflow illustrates the repeated transfer process to reach steady state (acclimation) and the subsequent dose-response exposure (0–1000 ppb) over 16 days. During the acclimation phase, short-chain fatty acid (SCFA) and gas yields were analyzed before each transfer (i.e., every 48 hours) until no significant variations were observed (i.e., microbial community reached steady state). All the culture was pooled at the end of this phase and used as the inoculum for the exposure phase. DNA was extracted from the pooled culture. During the exposure phase, SCFA, headspace gas, and microbial DNA were collected at Days 2, 8 and 16.

The bioreactor medium was prepared based on a published protocol (McDonald et al., 2013), with adjustments made to accommodate the bioreactor design. The medium was composed of the following (per liter): 0.0132 g CaCl2·2H2O, 0.1267 g NaCl, 0.04 g K2HPO4, 0.04 g KH2PO4, 0.0205 g MgSO4·7H2O, 0.001 g menadione, 0.0533 g peptone, 0.08 g yeast extract, 0.08 g pectin (from citrus), 0.08 g xylan (from corn core), 0.08 g arabinogalactan, 0.2 g starch (from corn), 0.12 g casein, 0.04 g inulin (from Dahlia tubers), 0.02 g bile salts, 0.16 g porcine gastric mucin (type II), 2 g NaHCO3, 0.005 g hemin, and 0.5 g L-cysteine HCl. The medium was boiled to facilitate the dissolution of ingredients under a continuous nitrogen sparge to remove dissolved oxygen. After cooling to room temperature, the medium was aliquoted into individual reactors, re-sparged with nitrogen, and sterilized via autoclaving (121°C, 20 minutes). For the exposure phase, 10 mL of the sterile medium was spiked with 5 µL of either the stock solutions (2, 0.2, or 0.02 µg/µL) or pure DMSO (vehicle control) using 10 µL syringes (Hamilton). This yielded final concentrations of 1000, 100, 10, and 0 ppb, respectively, with a final DMSO concentration of 0.05% (v/v).

### 2.4 Analytical methods

Bioreactor headspace gas pressure was measured using a barometer (General Tools). Methane (CH_4_), carbon dioxide (CO_2_), and hydrogen gas (H_2_) were quantified using a gas chromatograph (GC; 8890 System, Agilent) equipped with a flame ionization detector and a thermal conductivity detector (TCD). Due to the time-intensive nature of GC analysis, gas composition was determined from the headspace of a single reactor within each triplicate during the exposure phase. Since headspace sampling for GC precludes stable total pressure measurements, pressure was recorded from the remaining two reactors. Molar gas production (mmol) was subsequently calculated with the Ideal Gas Law, assuming a uniform gas composition across the triplicate set. Supernatant SCFAs, including acetic, propionic, and butyric acids, were quantified using a liquid chromatograph (1260 Infinity II Quaternary LC System, Agilent) equipped with an Aminex HPX-87H column (Bio-Rad).

### 2.5 Microbiome analysis

Total genomic DNA was extracted using the Qiagen DNeasy PowerSoil Pro Kit (QIAGEN) according to the manufacturer’s instructions. Full-length 16S rRNA gene amplicons were generated with the forward primer AGRGTTYGATYMTGGCTCAG and reverse primer AAGTCGTAACAAGGTARCY. High-fidelity (HiFi) sequencing was performed on the Sequel II system (PacBio). Raw reads were initially quality-filtered and primer-trimmed using the pb-16S-nf pipeline (version 0.6, available at https://github.com/PacificBiosciences/HiFi-16S-workflow) with default settings. The resulting reads were imported into R for filtering (minLen = 1000, maxLen = 1600, rm.phix = FALSE, maxEE = 2, other settings as default), denoising (DETECT_SINGLETONS = TRUE, other settings as default), and chimera identification and removal (minFoldParentOverAbundance = 3.5, other settings as default), using the R package dada2 (version 1.26.0) (Callahan et al., 2016). The processed reads were then imported to QIIME 2 (version 2024.10) (Bolyen et al., 2019) and mapped to the GTDB 220.0 database for taxonomic assignment. A custom Naive Bayes classifier was trained using the forward and reverse primers employed in this study. The relative abundances of microorganisms in each sample were exported at the species level, and species with low relative abundances (< 2%) were grouped as “others”. The feature table and taxonomy from QIIME2 and metadata were imported into R as a phyloseq object using the qiime2R package (version 0.99.6) (Bisanz, 2018), and then rarified using the phyloseq package (version 1.56.0) (McMurdie and Holmes, 2013) for alpha and beta diversity analyses. The rarefaction depth was 6,938, which was the size of the smallest library of all samples. Alpha diversity metrics, including Observed Amplicon Sequence Variants (Observed ASVs), Shannon Diversity, and Pielou’s Evenness, were calculated using the microbiome package (version 1.34.0) (Lahti and Shetty, 2012). The beta diversity matrix, Bray–Curtis dissimilarity, was calculated using the rbiom package (version 3.1.0) (Smith, 2025).

### 2.6 Statistics

All statistical analyses were conducted in R. To examine associations between Bray–Curtis dissimilarity and parameters of interest (e.g., DMSO vs. No-DMSO), permutational analyses of variance were conducted with the “adonis2” function (permutations = 9999, other settings as default) in the vegan package (version 2.7–2) (Oksanen et al., 2025). The homogeneity of group dispersions was assessed with the vegan “betadisper” and “permutest” functions (permutations = 9999, pairwise = TRUE, other settings as default). Differential abundance analysis was performed using the “ancombc2” function in the ANCOMBC package (version 2.10.1) (Lin and Peddada, 2024). Only ASVs with a prevalence of > 20% and a detection threshold of one were included in this analysis. *P*values were corrected using the default Holm method, and an adjusted *P*value (*q* value) of < 0.05 was considered statistically significant.

To evaluate changes in SCFA concentrations, gas production, and alpha diversity, linear mixed-effects models were employed using the “lmer” function in the R package lme4 (Bates et al., 2015), with Time, Concentration, and their interaction as fixed effects, and each experimental replicate as a random effect to account for repeated measures. All the data visualization was performed using the ggplot2 package (version 4.0.3) (Wickham, 2016).

## 3. Results

### 3.1 DMSO significantly altered SCFA and gas production, whereas AFB1 and FB1 showed limited effect

We first quantified the three major short-chain fatty acids (SCFAs) usually produced in the human gut—acetic, propionic, and butyric acids—across all experimental conditions, encompassing two xenobiotic types (aflatoxin B1 [AFB1] and fumonisin B1 [FB1]), five conditions (no-dimethyl sulfoxide [No-DMSO] control, 0, 10, 100, and 1000 ppb), and three time points (Days 2, 8, and 16) (**Figure 2**). To evaluate the influence of concentration and temporal progression on these metabolic profiles, we employed a linear mixed-effects model on SCFA levels for each mycotoxin (**Table S1**). Our analysis revealed that while the two mycotoxins might exert an acute initial effect, the DMSO vehicle emerged as the primary driver over time. Upon the initiation of AFB1 exposure (Day 2), acetic acid levels in the 1000 ppb group were significantly elevated compared to the 0-ppb group (*p*= 0.016). However, this AFB1-specific modulation was transient and did not persist as the duration of exposure increased. Starting at Day 8 and persisting through Day 16, no statistical difference was observed between any of the AFB1-treated groups (i.e., 10, 100, 1000 ppb) and the 0-ppb group across all analyzed acids (*p*s > 0.05). On the contrary, DMSO in DMSO-containing groups (i.e., 0, 10, 100, 1000 ppb) showed no significant deviation from the No-DMSO control at Day 2 (*p*s > 0.05), but at later time points drove a significant increase in acetic acid and a substantial decrease in propionic acid levels relative to the No-DMSO control (*p*s < 0.05). Moreover, DMSO reduced butyric acid production, showing significant (*p*s < 0.05) or near-significant (0.053 < *p*s < 0.13) deviations from the No-DMSO control at Days 8 and 16.

**Figure 2.**
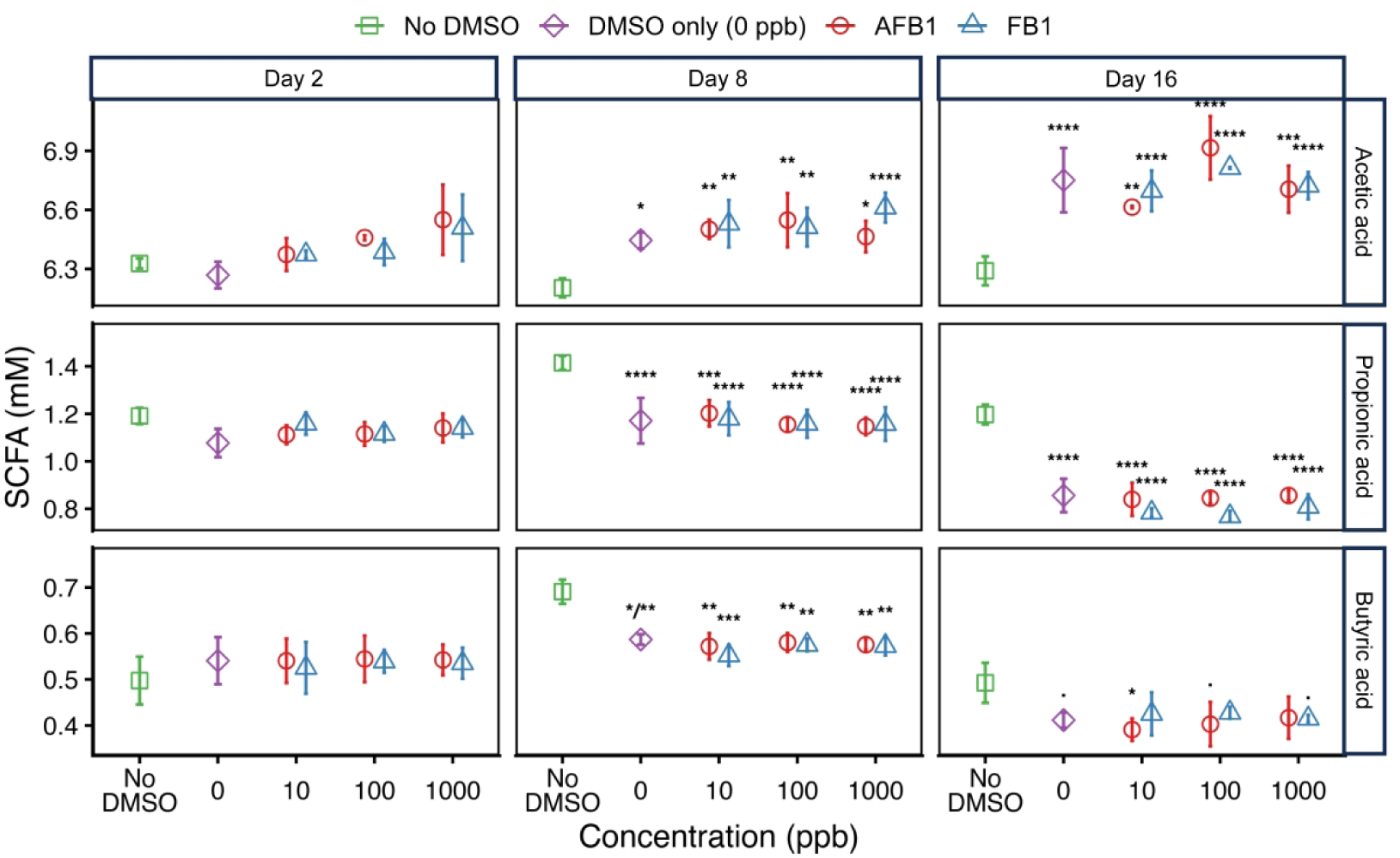
Short-chain fatty acid (SCFA) levels under varying concentrations of aflatoxin B1 (AFB1) and fumonisin B1 (FB1), stratified by time and SCFA type. Note that the no- dimethyl sulfoxide (No-DMSO) and 0-ppb groups are identical between the AFB1 and FB1 datasets. Error bars denote one standard deviation from biological triplicates (*n* = 3). Asterisks represent significant differences between each treatment group and the No-DMSO control, as determined by linear mixed-effects models (•: *p*= 0.05; *: *p*< 0.05; **: *p*< 0.01; ***: *p*< 0.001; ****: *p*< 0.0001; all Holm corrected). Note that for butyric acid at Day 8, the comparison between the 0 ppb and the No-DMSO control reached different significance thresholds in the independently fitted models (AFB1 model: *p*< 0.05; FB1 model: *p*< 0.01).

The FB1 results followed a similar temporal pattern. At Day 2, the 1000-ppb group elicited a significant initial elevation in acetic acid concentrations compared to the 0-ppb group (*p*= 0.026). As seen with AFB1, each DMSO-containing group at this early stage did not differ significantly from the No-DMSO control (*p*s > 0.17). By Day 16, DMSO induced a significant increase in acetic acid (*p*s < 0.0001) levels, a profound decline in propionic acid (*p*s < 0.0001), and a marginal decrease in butyric acid levels (0.058 < *p*s < 0.17).

In addition to SCFA profiles, we also analyzed gas production across all experimental conditions. Parallel to the acid results, gas levels were significantly influenced by the DMSO vehicle employed for mycotoxin delivery (**Figure S1 and Table S2**). For AFB1, gas yields were immediately dominated by DMSO at Day 2, which induced a significant suppression of methane (CH4; *p*s < 0.0001) and a simultaneous increase in carbon dioxide (CO_2_) levels (0.025 < *p*s < 0.72) in DMSO-containing groups, relative to the No-DMSO control. This solvent-driven divergence persisted through Day 16, where CH4 levels remained markedly lower than the control (*p*s < 0.0001). The low-concentration (10 ppb) AFB1 level significantly altered CH4 yields relative to 0, 100 and 1000 ppb concentrations (*p*s < 0.013) at Day 2, but this acute impact was transient and became statistically undetectable after Day 8 (*p*s > 0.05).

Compared with AFB1, FB1 showed a more pronounced acute response. At Day 2, the 10-, 100-and 1000-ppb groups significantly inhibited CH4 generation compared to the 0-ppb group (*p*s < 0.05), but this mycotoxin-induced effect was specific to the initial stage of exposure and disappeared by later time points. Collectively, these findings indicate that for both mycotoxins, the long-term metabolic shifts were driven by the DMSO solvent rather than the mycotoxin itself.

### 3.2 Microbial composition was significantly impacted by DMSO, not by mycotoxins

Pairwise comparisons between individual mycotoxin concentrations revealed no significant differences in microbial community composition (*p*s > 0.05), indicating a lack of dose-dependent responses. However, a significant global shift was observed when comparing No-DMSO controls to pooled DMSO-treated groups (0–1000 ppb) (**Figure 3**). While this divergence was not yet significant at Day 2 (AFB1: *p* = 0.22; FB1: *p* = 0.071), it became increasingly pronounced starting at Day 8 and continued through Day 16 (*p*s < 0.005 for all comparisons). Notably, this observed structural divergence was accompanied by significant differences in group dispersion (*p*s < 0.05), suggesting that increased community stochasticity within the treated groups might contribute to the separation. In contrast, treatment effects on alpha diversity were minimal; significant differences were restricted to Day 8, where only the Observed Amplicon Sequence Variants (Observed ASVs) and Shannon Diversity index distinguished the 0-ppb group from the No-DMSO control (**Figure S2, Table S3**).

**Figure 3.**
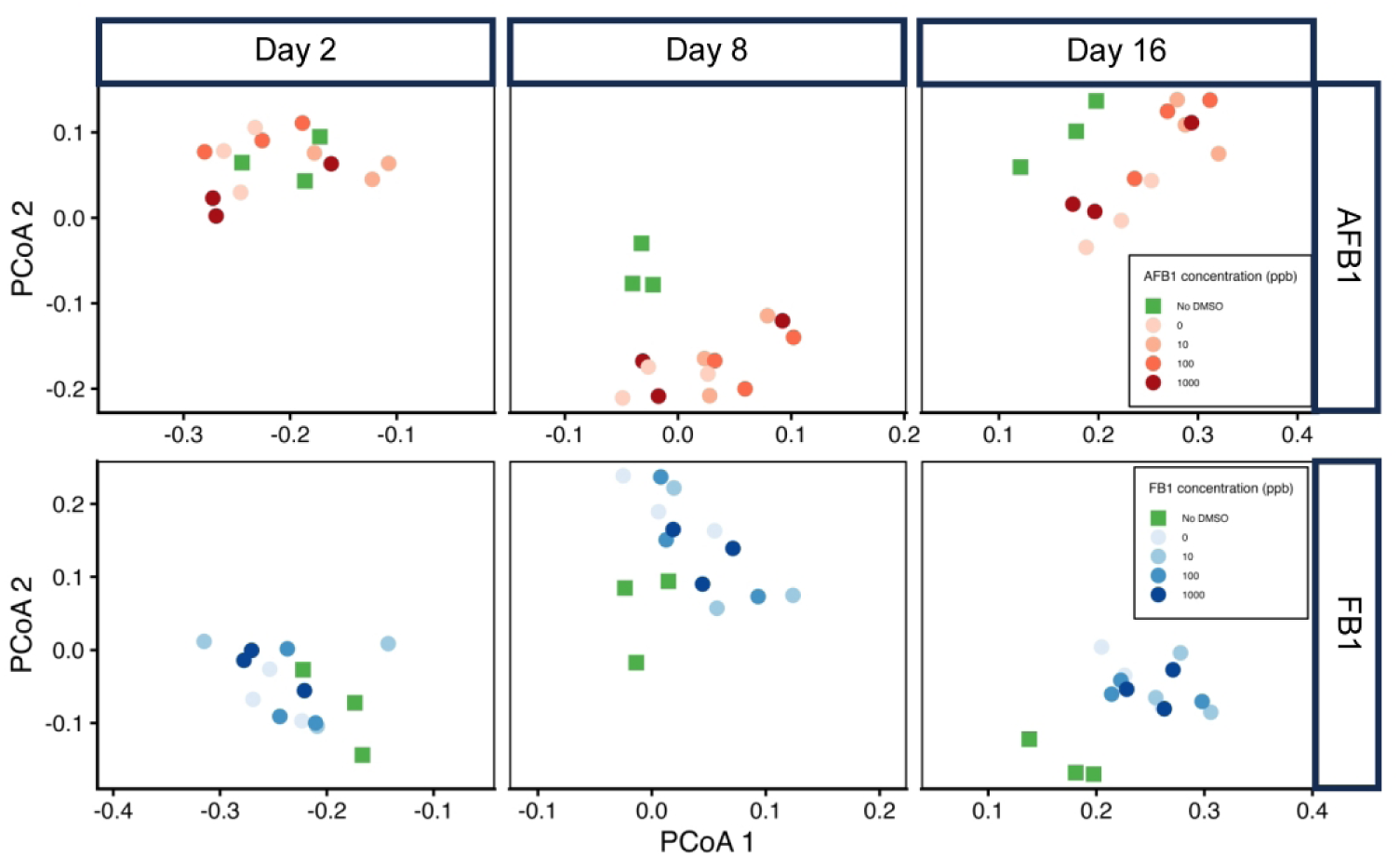
Beta diversity plot showing the Bray-Curtis dissimilarity, stratified by time and mycotoxin type. Note that the no-dimethyl sulfoxide (No-DMSO) and 0-ppb groups are identical between the aflatoxin B1 (AFB1) and fumonisin B1 (FB1) datasets.

The overall taxonomic composition remained largely conserved in DMSO-treated groups at the species level, regardless of the mycotoxin types and concentrations (**Figure S3**). Differential abundance analysis further characterized the microbial response to the solvent vehicle and mycotoxin treatments. Despite global community stability at Day 2, a single ASV belonging to *Desulfovibrio desulfuricans* was significantly enriched in pooled DMSO-treated groups relative to No-DMSO controls (AFB1: *q* = 0.016; FB1: *q* = 0.00019; **Figure S4**). This targeted enrichment was consistent across both mycotoxin types and persisted through Day 16. Specifically, while no individual mycotoxin concentration (0, 10, 100, or 1000 ppb) differed significantly from the No-DMSO control at Day 2, a robust enrichment of the same *D. desulfuricans* ASV emerged across each of these concentrations relative to the No-DMSO control starting at Day 8, reaching relative abundances of 6.0-11% (**Figure 4**). Notably, these treated groups remained indistinguishable from one another, as no ASVs were differentially abundant between any two mycotoxin concentrations (*q*s > 0.05). This indicates that the observed shifts were primarily a response to the DMSO vehicle rather than a dose-dependent reaction to the mycotoxins.

**Figure 4.**
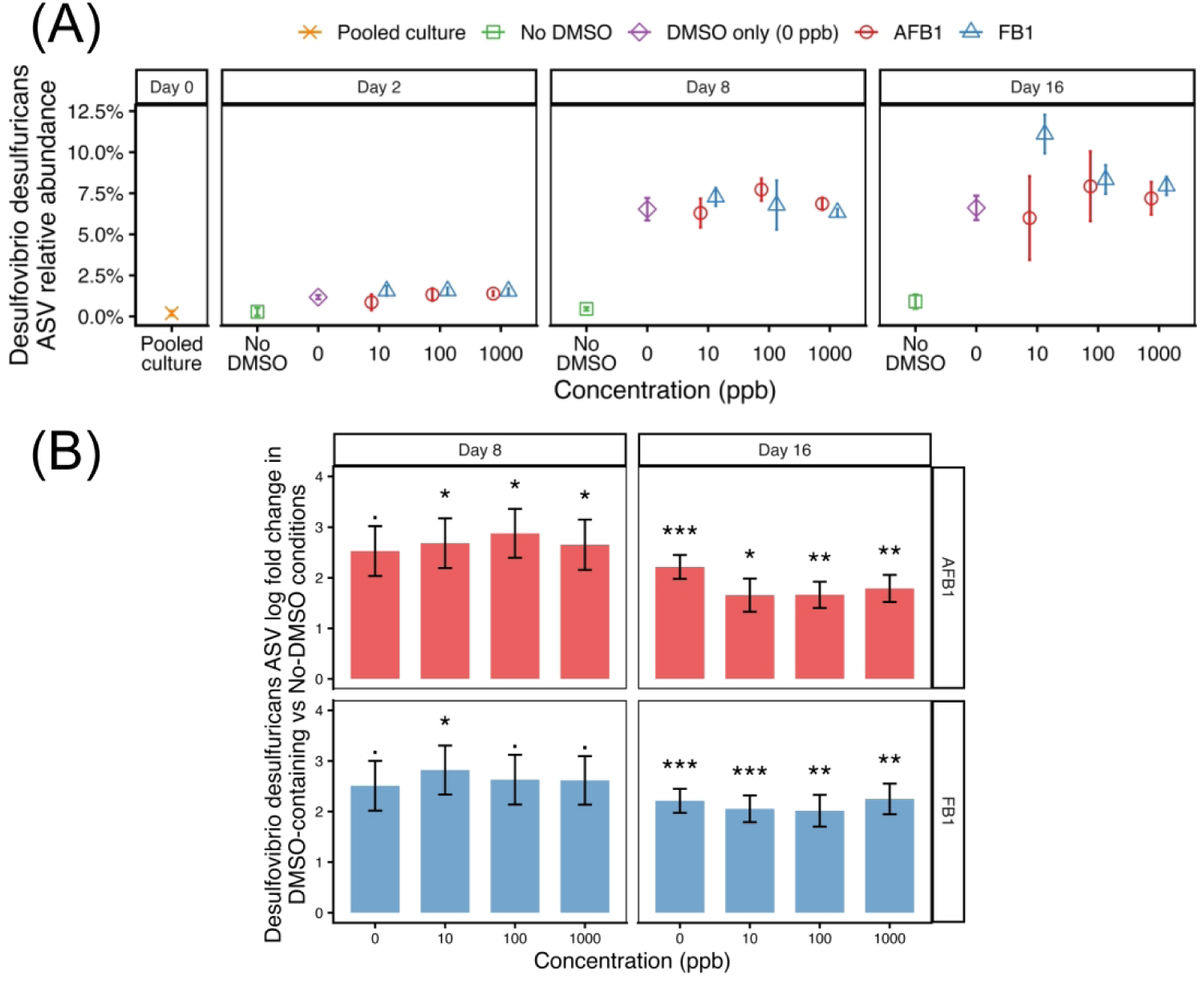
(A) Relative abundances of a *Desulfovibrio desulfuricans* amplicon sequence variant (ASV) in the pooled culture, and under varying concentrations of aflatoxin B1 (AFB1) and fumonisin B1 (FB1) stratified by time. Note that the no-dimethyl sulfoxide (No-DMSO, and 0-ppb groups are identical between the AFB1 and FB1 datasets. Error bars denote one standard deviation from biological triplicates (*n* = 3). (B) Differential abundance analysis showing the enrichment of the *D. desulfuricans* ASV in DMSO-containing reactors compared with No-DMSO controls. Bars represent the log fold change of ASVs significantly associated with AFB1 (red) and FB1 (blue) treatments. Error bars denote one standard error. •: *p*= 0.05; *: *p*< 0.05; **: *p*< 0.01; ***: *p*< 0.001 (all Holm corrected).

Moreover, a few other ASVs were identified as differentially abundant between pooled DMSO-treated groups and No-DMSO controls. One ASV belonging to *Parabacteroides goldsteinii* was significantly lower in abundance in the DMSO-treated group at Day 8, and another ASV belonging to *Phocaeicoladorei* was significantly higher in abundance at Day 16 (**Figure S4**).

Overall, the microbial community dynamics were dominated by the solvent effect, which masked any potential dose-dependent responses to the mycotoxins. The structural divergence across the community was largely attributable to DMSO, which triggered a targeted bloom of *D. desulfuricans*.

## 4. Discussion

A central finding of this study was the influence of the dimethyl sulfoxide (DMSO) solvent on the gut microbiota, which effectively masked potential dose-dependent responses to aflatoxin B1 (AFB1) and fumonisin B1 (FB1). At Day 2, the highest concentrations appeared to influence short-chain fatty acid (SCFA) production, indicating that initial microbial response might be impacted by mycotoxins themselves, not by DMSO. However, as the reaction progressed to Day 8 and beyond, DMSO respiration became more dominated and overwhelmed the mycotoxin trends. While DMSO is a solvent frequently used for studying interactions between compounds of interest (e.g., toxins, drugs, food compounds, and microbial metabolites) and gut microbiota (Chen et al., 2018; Lai et al., 2018; O’Connor et al., 2019; Tang et al., 2021; Osei-Owusu et al., 2022; Li et al., 2024; Müller et al., 2024; Tian et al., 2024; Yang et al., 2024; Roux et al., 2025; Wang et al., 2025), the majority of past research omitted a No-DMSO control and a comparison of treatments against a DMSO-solvent baseline. Even when such controls are included, the observed impact can vary across different experimental setups. For instance, a recent study evaluating 1% DMSO on honeybee gut microbiota reported no significant differences between solvent-treated and no-solvent groups after 10-day exposure (Wang et al., 2025). In another study on the impact of triclosan on zebrafish, comparisons between 0.01% DMSO and No-DMSO controls revealed shifts in microbial relative abundances after 120 days of exposure; however, the potential role of the DMSO vehicle in these shifts remained largely unexplored (Tang et al., 2021). One plausible explanation for these discrepancies is that the gut microbiota in previous studies may have lacked specific DMSO-metabolizing species, allowing the community to remain stable. In our study, although *Desulfovibrio desulfuricans* was present at trace levels (0.19%) in the pooled culture, it became the most enriched taxon across all DMSO-treated replicates within eight days. This rapid emergence highlights the profound selective pressure exerted by the solvent. Therefore, a no-solvent control is necessary to avoid misattributing solvent-driven shifts to the compounds under investigation. Alternatively, using endogenous food components as solvents (e.g., peanut oil for mice on a peanut-supplemented diet) might help minimize non-target vehicle impacts (Dean et al., 2025).

The targeted enrichment of *D. desulfuricans* across all DMSO-treated groups suggests a specific adaptation. *Desulfovibrio* species are known for their sulfate-reducing capabilities, including the ability to utilize DMSO as a terminal electron acceptor and reduce it to dimethyl sulfide (DMS) under anaerobic conditions (Jonkers et al., 1996). This metabolic shift is further supported by the significant inhibition of methane (CH_4_) production across all toxin concentrations (0–1000 ppb). Since methanogenesis requires electrons for carbon dioxide (CO_2_) reduction, the presence of DMSO likely created a competitive electron sink where electrons were prioritized for DMSO reduction by *Desulfovibrio* rather than for methanogenesis. Nevertheless, the amount of DMSO supplemented in our experiment (5 μL per reactor, which could theoretically accept ∼141 μmol of electrons and lead to an ∼18 μmol loss in CH_4_) could not fully account for the observed CH4 production differences between DMSO-containing and No-DMSO conditions (> 40 μmol), based on electrons. This indicates that methanogenesis might be inhibited by other factors. It should be noted that a solid electron balance calculation might be limited by the use of a single gas composition value across replicates in this study. Future research utilizing high-resolution, replicate-specific gas measurements is needed to precisely trace the electron flow and confirm the bioenergetic advantage provided by DMSO respiration.

*Desulfovibrio* can contribute to intestinal pathogenesis through the production of hydrogen sulfide, which is cytotoxic to colonocytes at elevated concentrations (Singh et al., 2023; Huang et al., 2024). *Desulfovibrio* can also release lipopolysaccharide that trigger the production of pro-inflammatory cytokines (Weglarz et al., 2003; Węglarz et al., 2007; Zhou et al., 2024). Furthermore, *Desulfovibrio* is able to affect the growth of mucin-degrading bacteria and compromise the integrity of the epithelial barrier (Zhang et al., 2023). Given that some studies investigating the toxicological profiles of AFB1 and FB1 rely on DMSO as a carrier solvent, it is plausible that the pro-inflammatory signatures frequently attributed to these toxins are a confounding result of vehicle-induced shifts in the gut microbiota. Future investigations into interactions between microbiota and mycotoxins (and other xenobiotics) must account for the independent effects of solvents to ensure that observed inflammatory outcomes are accurately interpreted.

Our study also suggests that the bioreactor system could be a complementary tool to *invivo* models for a quick solvent sensitivity check. While solvents are often assumed to be inert at low concentrations, our data challenges this assumption in a direct-exposure context. The lack of significant solvent effects observed in previous *in vivo* studies likely results from host physiological processes (e.g., rapid absorption or hepatic clearance) buffering the gut ecosystem. However, relying solely on this host-mediated protection may introduce unrecognized confounding variables. The bioreactor environment circumvents this host interference, unmasking the intrinsic vulnerability of the gut microbiota to common solvents.

Overall, our findings show that DMSO as the solvent vehicle can have unintended effects on gut microbiota composition and function. While these effects were minimal on Day 2, they became increasingly pronounced by Day 8. This highlights a clinical need to validate any solvent used in xenobiotic exposure studies to ensure it does not accidentally disrupt microbial balance. However, this study has several limitations. First, a dose-response evaluation for DMSO was not conducted, leaving the threshold for a definitively inert vehicle concentration unidentified for this specific microbial community. Second, while pooling donor samples allowed for a standardized and reproducible microbial community across replicates, this approach minimizes inter-individual variability and may not reflect the diverse susceptibility found in broader populations. Additionally, since the biotransformation of the tested compounds was not monitored in this study, it remains unclear whether the observed microbial shifts were impacted by the parent xenobiotics, their degradation products, or the modulation of specific taxa capable of compound sequestration or degradation, which would further impact the overall community response. Finally, while the 16-day experimental design identified microbial shifts, it may not fully capture the consequences of chronic exposure, which in human populations typically spans years or decades. Future research should integrate biotransformation of xenobiotics in long-term models and rigorous vehicle validation to more accurately characterize their chronic impacts on the human gut microbiota. Moreover, conducting studies on individual microbial donors will be essential to understanding personalized reactions to xenobiotics.

## Data availability statement

The raw 16S rRNA gene amplicon data were deposited in the National Center for Biotechnology Information (NCBI) Sequence Read Archive (SRA) database (BioProject accession number: PRJNA1510332).

## Ethics statement

Human fecal sample collection for the bioreactor study was approved by Arizona State University Institutional Review Board (STUDY00004850). The study was conducted in accordance with the local legislation and institutional requirements.

## Author contributions

QC: Conceptualization, Data curation, Formal analysis, Funding acquisition, Investigation, Methodology, Software, Visualization, Writing – original draft, Writing – review & editing. HG: Conceptualization, Data curation, Investigation, Methodology, Writing – review & editing. ASC: Conceptualization, Data curation, Investigation, Methodology, Writing – review & editing. LEV-G: Conceptualization, Funding acquisition, Methodology, Project administration, Supervision, Writing – review & editing. RKB: Conceptualization, Funding acquisition, Methodology, Project administration, Resources, Supervision, Writing – review & editing.

## Acknowledgments

The authors would like to thank Dr. Peter Rohloff, Dr. Ann Miller, Dr. Gabriela Montenegro, and Olga Torres for advice in experimental design and for collaborative grant application.

