## Supplementary figures for "Reconsidering the Use of Dimethyl Sulfoxide for Xenobiotic-Gut Microbiota Interaction Studies"

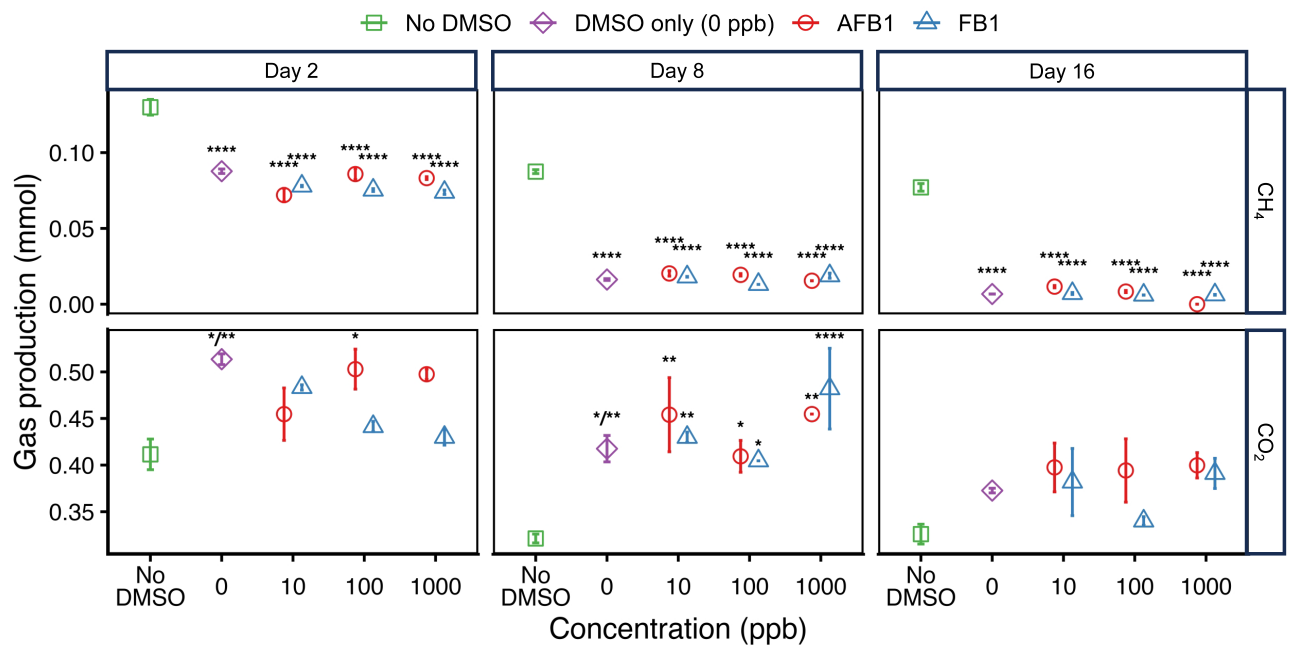

**Figure S1** Methane (CH_4_) and carbon dioxide (CO_2_) production under varying concentrations of aflatoxin B1 (AFB1) and fumonisin B1 (FB1), stratified by time and gas type. The no-dimethyl sulfoxide (No-DMSO) and 0-ppb groups are identical between the AFB1 and FB1 datasets. For each experimental condition, the molar quantity of individual gas components (in mmol) was calculated by applying the Ideal Gas Law to the total gas volume (measured in duplicate, *n* = 2) and the fractional composition (measured from a third biological replicate, *n* = 1). Error bars denote the range of calculated molar quantities (*n* = 2). Asterisks represent significant differences between each treatment group and the No-DMSO control, as determined by linear mixed-effects models (•: *p* = 0.05; *: *p* < 0.05; **: *p* < 0.01; ***: *p* < 0.001; ****: *p* < 0.0001; all Holm corrected).

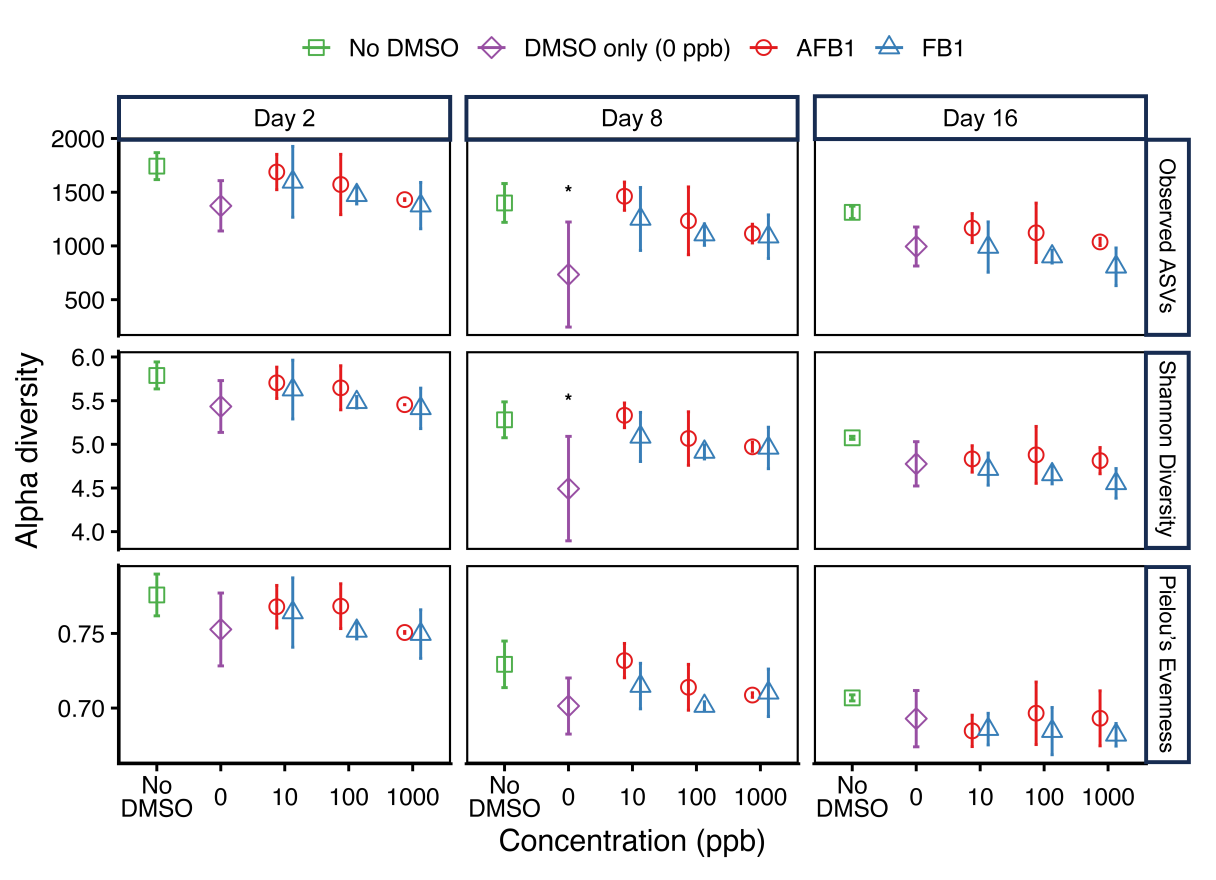

**Figure S2** Alpha diversity metrics under varying concentrations of aflatoxin B1 (AFB1) and fumonisin B1 (FB1), stratified by time and metric type. Note that the no-dimethyl sulfoxide (No-DMSO) and 0-ppb groups are identical between the AFB1 and FB1 datasets. Error bars denote one standard deviation from biological triplicates (*n* = 3). Asterisks represent significant differences between each treatment group and the No-DMSO control, as determined by linear mixed-effects models (*: *p* < 0.05; Holm corrected).

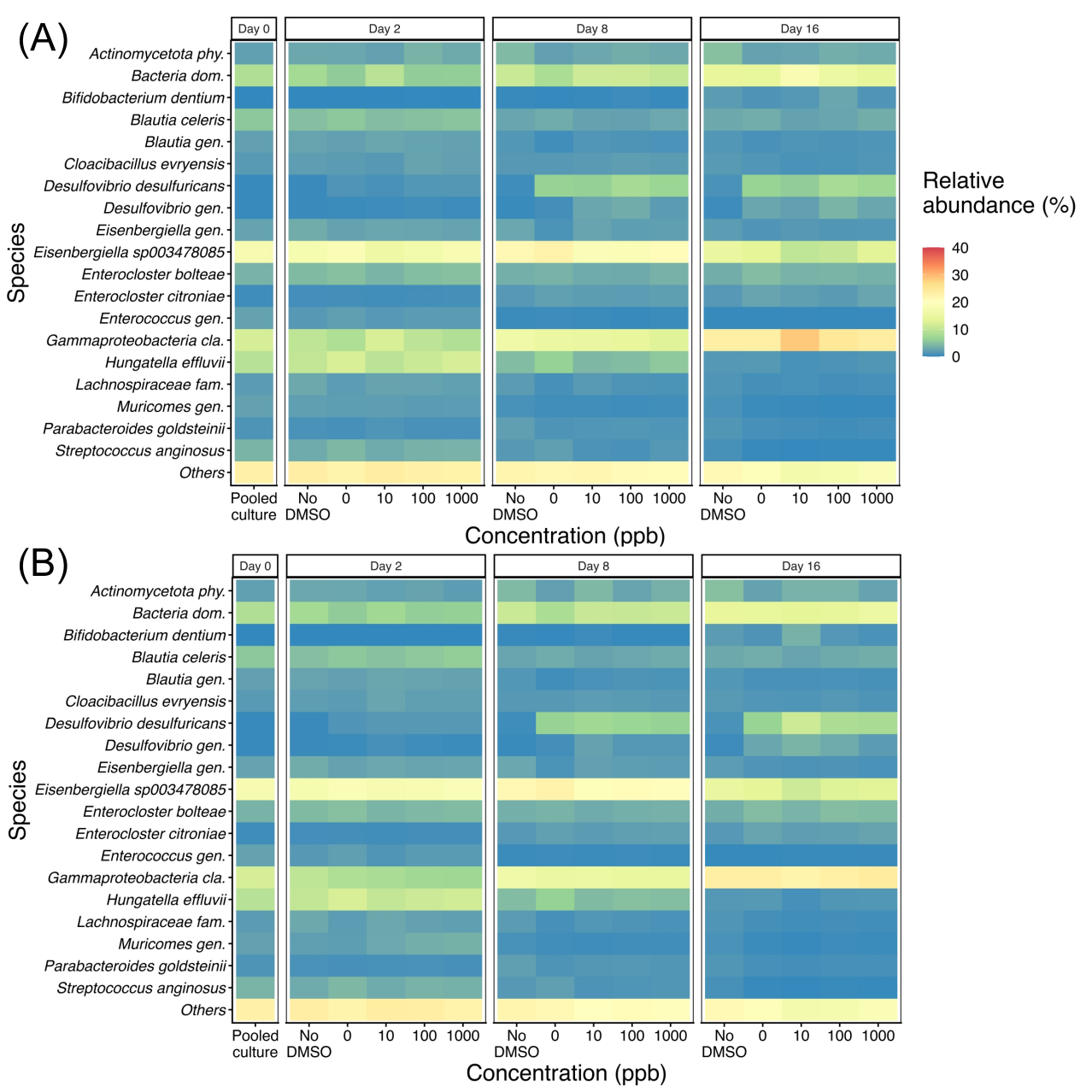

**Figure S3** Heatmaps displaying the relative abundance of species that exceeded 2% in at least one sample across (A) aflatoxin B1 (AFB1) and (B) fumonisin B1 (FB1) treatment groups, with data stratified by sampling time. Note that the no-dimethyl sulfoxide (No-DMSO) and 0-ppb groups are identical between the AFB1 and FB1 datasets.

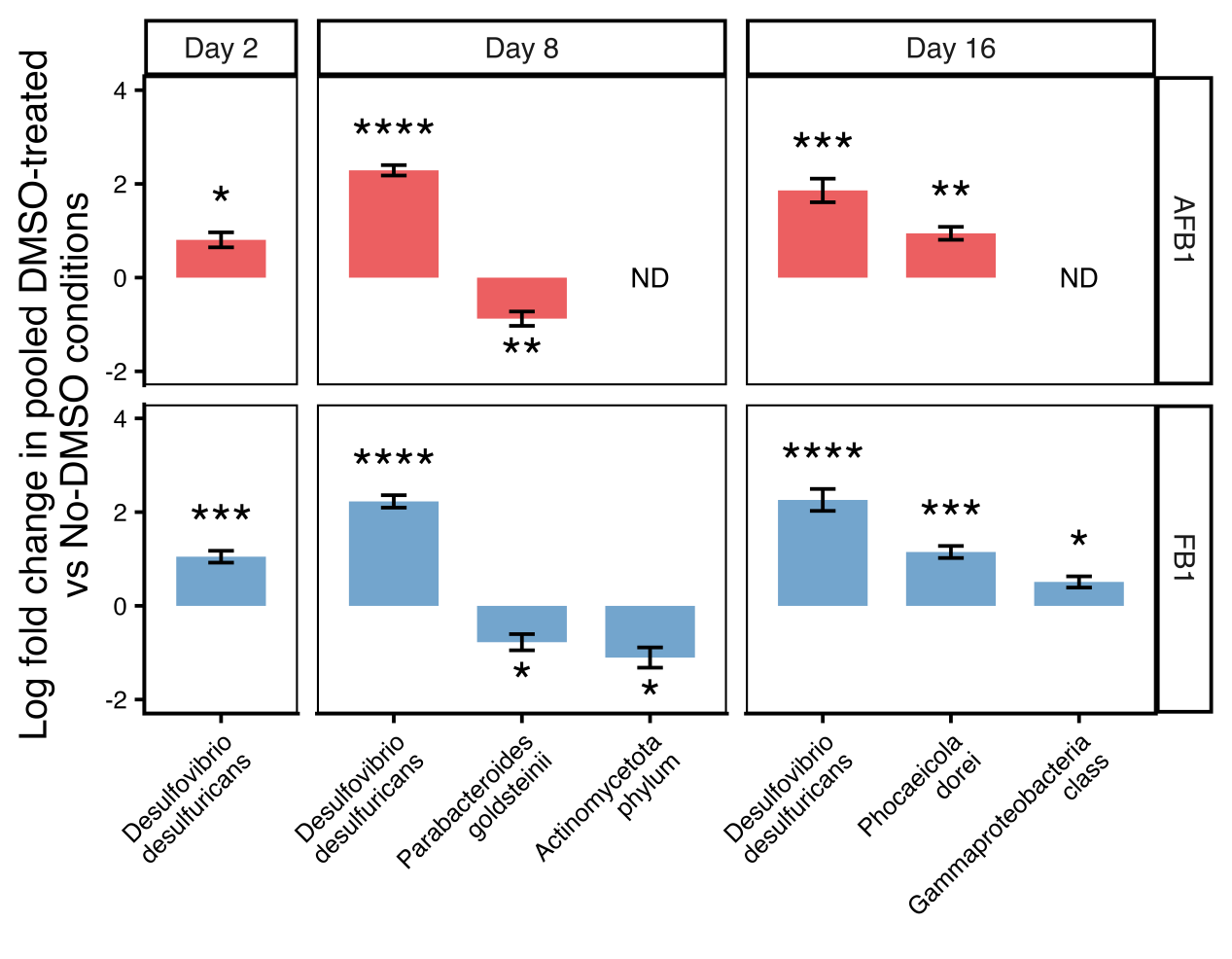

**Figure S4** Differential abundance of bacterial amplicon sequence variants (ASVs) in dimethyl sulfoxide (DMSO)-treated reactors relative to No-DMSO controls. Bars represent the log fold change of ASVs significantly associated with aflatoxin B1 (AFB1, red) and fumonisin B1 (FB1, blue) treatments (pooled data spanning concentrations 0-1000 ppb for each mycotoxin). Positive values indicate ASV enrichment in treatment groups, while negative values indicate ASV abundance decrease. Error bars represent one standard error. ND: not detected; *: *p* < 0.05; **: *p* < 0.01; ***: *p* < 0.001; ****: *p* < 0.0001 (all Holm corrected).
